# ORTHOTOPIC TUMOUR INDUCTION PROTOCOL SHAPES TUMOUR MICROENVIRONMENT

**DOI:** 10.64898/2026.10.31.763307

**Authors:** Shauna McClelland, Yanbing Liu, Gloria Allocca, Nikita Lewis, Amanda Eakin, Cristina M Branco

## Abstract

Robust preclinical cancer models are essential for studying tumour progression and evaluating therapeutic strategies. Orthotopic implantation of tumour cells into the mammary gland is widely used to model triple-negative breast cancer (TNBC), yet substantial variation exists in the number of cells used to initiate tumours, often without explicit consideration of how this parameter shapes tumour biology. Here, we systematically assess how tumour cell number at implantation influences the tumour microenvironment using the 4T1 orthotopic TNBC model. Mammary tumours initiated with different cell numbers exhibited comparable early growth kinetics and gross size, yet diverged markedly in cellular density, hypoxia, necrosis, proliferation, vascular integrity, and immune cell infiltration. High-density tumours displayed increased HIF-1α staining and necrosis, alongside differences in vascular-associated markers and increased macrophage recruitment. Conversely, low-density tumours remained highly proliferative, less necrotic and exhibited higher PDGFRβ staining. Importantly, tumour cells disseminated systemically within days of implantation across all conditions, although differences in colony recovery and *ex vivo* expansion were not statistically significant. Together, these findings demonstrate that tumour initiating protocols are powerful determinants of tumour state, with potential implications for dissemination frequency and tumour cell fate at distant sites. Rather than representing a source of experimental variability alone, this understanding provides an opportunity to intentionally tailor orthotopic models to investigate specific stages of disease progression and therapeutic response, thereby enhancing cross-study reproducibility and translational relevance.

**Summary Statement:** Orthotopic breast tumours generated with different cell numbers display distinct microenvironments, demonstrating that tumour induction protocols critically shape experimental outcomes and translational interpretation.

## Introduction

The impact of research in cancer biology, progression and dissemination relies on robust experimental modelling to enable its translational value for human disease. Over decades, researchers have improved and refined *in vitro* and *in vivo* model systems, to study and validate molecular and physiological parameters that may lead to identification of therapeutic targets or better treatment approaches. Mouse models are overwhelmingly used for preclinical studies of many cancer types, owing to pathological properties that, to a comfortable extent, mirror human disease, as well as being relatively easy to manipulate genetically, with life cycles shorter than other mammalian models, allowing generation of genetically engineered mouse models (GEMM) with relative efficiency.

The evolution of mouse models for the study of cancer spans diverse strategies. Heterotopic implantations (e.g., subcutaneous models) are useful in studying cancer cells within a tissue environment as opposed to in isolation, are an adequate option to study cancers difficult to implant orthotopically or for which suitable genetically-induced models do not yet exist, and an excellent starting point for preclinical drug testing(Stribbling & Ryan, 2022). With the established realization that the tumour microenvironment influences tumour cell behaviour, these models provide limited value in representing the biology of human disease and forecasting its progression in a real-life scenario(Devaud et al., 2014; Fernandez et al., 2023). Orthotopic models, in which tumour cells or tumour fragments are placed into the sites in which the tumours would naturally develop, offer the advantage of providing the appropriate tissue context. However, to investigate behaviours of human cancer cells (xenografts), these can only graft in the absence of a competent immune system, a substantial caveat in reflecting its fundamental contribution to tumour growth and metastatic processes(Biray Avci et al., 2024; Fernandez et al., 2023; Guo et al., 2024; Zhang et al., 2025; Zhou & Zhou, 2025). Circumventing this critical aspect of cancer growth will also compromise the ability to develop and predict their response to different therapeutic approaches, which heavily hinge on systemic, multicellular responses. This approach not only creates artifacts in our interpretations of the biology but also limits applications in the clinic. Elaborate GEMM have significantly improved the experimental landscape for cancers with known genetic drivers such as KRAS in lung(Drosten et al., 2018), pancreatic(Lee et al., 2016; Mazur & Siveke, 2011), or colorectal cancers, PTEN and tkp53 in prostate cancers (Arriaga & Abate-Shen, 2019; Kaushal et al., 2024; Mejía-Hernández et al., 2022), and often breast cancers (Regua et al., 2021; Rennhack et al., 2019). These can be manipulated to induce tumours in the right tissues, in a fashion that much more closely resembles what occurs in human cancer pathophysiology and continue to offer strong opportunities for intervention. However, they too come with limitations: in TNBC, there is high heterogeneity and processes that cannot be fully recapitulated, and increasing the genetic complexity of these models inevitably incurs undesirable and unrelated effects (Doha et al., 2023; Grasset et al., 2022). This is the breast cancer subtype with the highest metastatic and mortality rate; approximately 5 % carry the BRCA1 or 2 mutations (bringing the relevance to ∼5 % of all breast cancer cases), and the large majority of patients with this diagnosis have no targetable or treatable molecular elements that can aid the approach to their therapy.

*In vivo* mouse models of TNBC are relatively reliable. These include orthotopic implantation of cells directly into mammary glands, which allows monitoring the development of one tumour, its dissemination and effects on tumour-free organs. The orthotopic route bypasses the confounding effects of genetically driven spontaneous models, such as the MMTV-PyMT, which will cause mice to asynchronously develop multiple tumours in distinct mammary glands. While each tumour can be observed in isolation, the systemic effects of tumour bearing cannot be accurately matched to tumour burden or staging, metastatic patterns, or windows for intervention.

Implantation of 4T1 tumour cells into the mammary gland is one of the most common orthotopic allograft models of TNBC, and has provided invaluable opportunities to study this disease. Unlike most murine cancer models which overwhelmingly metastasise to the lung, 4T1 tumours disseminate systemically, and this has provided foundational insights into molecular mechanisms of organ specific colonization(Pulaski & Ostrand-Rosenberg, 2001). It also has an intrinsic resistance to 6-Thioguanine(Liu et al., 2019; Paschall & Liu, 2016), allowing retrieval of cells from multiple organs, and selectively expand these cells to characterise their properties as a function of site, stage or treatment challenges. While it has increasingly been favoured over other similar allografts, there is little consensus across the literature as to how this model should be established. The wide range of experimental set-ups within this model alone, and tumours are generated in mice between 4- and 10-weeks old, and using as few as 5 × 10^2^ cells (Gregório et al., 2016), through to 5 × 10^5^ (Zhang et al., 2023), 1 × 10^6^ cells (Tao et al., 2008) or even as many as 2 × 10^6^ cells per mammary gland (Li et al., 2021) with an equally broad range of outcomes. One important aspect is indeed the heterogeneous response to therapy, as reported in (Gregório et al., 2016) which is critically important for TNBC prognosis and patient management in the absence of targeted and efficacious treatment options for this patient group.

While artificial syngeneic models require that enough cells are implanted to ensure tumour take, in reality tumours are not established at a rate of thousands of cells at a time. The microenvironments of these artificially implanted tumours will evolve impacted by the protocol used to generate it, such as the number of cells. These unique microenvironments will influence dissemination rates, possibly also dissemination patterns, or response to treatments.

In this study, we generated primary 4T1 tumours by implanting 1 × 10^4^, 5 × 10^4^ or 2.5 × 10^5^ cells into one mammary gland, and examine how this aspect of the tumour initiation protocol contributes to the tumour microenvironment over time.

## Materials and methods

### Animal Experiments

Animal studies were conducted under project license PPL2859, approved by the NI Department of Health/ United Kingdom Home Office and the institutional ethical committee, and compliant with the ethical principles of the Animals (Scientific Procedures) Act 1986. BALB/c female mice aged 8-weeks received one orthotopic injection into the mammary fat pad of 1 × 10^4^, 5 × 10^4^, or 2.5 × 10^5^ 4T1^GFP+Luc^ cells or saline sham implant (n = 3 per experimental group) grown at 37 °C, 5 % O_2_ and 5 % CO_2_. Primary tumours were measured trice weekly with callipers. One experimental group was terminated at day 3 post implantation (early onset, presumed pre-metastatic), to assess parameters of early tumour formation, a second group at day 21 (established tumours and potentially early metastasis) and a third group 35 days post implantation, to represent advanced and likely disseminated disease. Primary tumours, contralateral mammary glands and visceral organs were collected in 10 % Formalin and fixed for up to 48 h at 4 °C, then embedded in paraffin (n=3/group/time point); blocks were sliced into 5-10 µm sections and mounted onto microscope slides for subsequent histological analyses. A separate experiment using the same implantation protocol and cell density groups was performed to investigate early tumour cell dissemination (n=4 mice per group). Here, brain, lung, liver (metastatic) and kidney (non-metastatic control) from tumour bearing mice were harvested 3 and 7 days after implantation to retrieve 4T1^GFP+Luc^ cells.

### Haematoxylin and Eosin (H&E) Staining

The slides were deparaffinised twice in Histoclear (Sigma, MKCG7250) for 5 min each and rehydrated by immersing into graded alcohols, in sequential 3 min incubations from 100 % ethanol twice, followed by one incubation each in 95 %, 70 %, and 50 % ethanol before a final 5 min wash in deionized water. Nuclei were stained using Haematoxylin (Sigma, 1051750500) for 3 min, washed in tap water until the water ran clear before being immersed in bluing agent (0.1 %, NaHCO_3_, pH8.0) for 30 s. Slides where immersed for 10 dips in 95 % EtOH before incubating with Eosin (cytoplasm stain, Sigma, 1098441000) with glacial acetic acid added 1:500, for 2 min, and then dipped once in tap water. Slides were dehydrated using increasing concentrations of ethanol (95 %, 100 %, 100 %, 100 %) for 10 dips in each, before being immersed in Histoclear twice, for 3 min each. The slides were left out to dry overnight and mounted with quick-drying mounting media (Eukitt, BCBT0137) or glycerol jelly (prepared in house, 5 g Gelatine, 50 mL diWater and 50 mL Glycerol). n = 3 mice per group, 4 sections per mouse, minimum 1 image per section (≥ 12 datapoints per experimental group).

### Immunohistochemistry (IHC)

Slides were deparaffinised and rehydrated as above. Next, antigen retrieval was performed using Citrate-based buffer (H-3300, Vector Labs), prepared according to manufacturer’s instructions (15 mL citrate in 1600 mL deionised water). Slides were submerged in a glass box, covered partially with a glass lid and microwaved on high power for 4 min, followed by a further 11 min at low power. After cooling at room temperature in the same buffer for 30 min, slides were washed 3 times in PBS, for 5 min each. To quench endogenous peroxidase, each section was incubated with 150 µl of 3 % H_2_O_2_ (Sigma 216763) in methanol for 10 min. Sections were subsequently blocked with Universal Protein Blocking Agent (GeneTex, GTX30963) for 7 min, rinsed quickly in PBS and blocked further with 5 % serum of the host of the secondary antibody for each target, when available. For biotinylated antibodies, a biotin/streptavidin block was prepared (Vector labs, SP2002). When primary antibodies were raised in mouse, an extra block of MOM blocking solution (Vector, MKB-2213) was performed for 1 h in addition to the standard blocking step. Blocking solutions were rinsed off 2-3 times, then washed for 5 min in PBS, and sections permeabilised with 0.4 % Triton X-100 (Sigma, T8787) in PBS for 10 min. Slides were washed for 5 min in PBS, and a hydrophobic barrier was drawn around each of the sections with a water-repellent pen. Primary and secondary antibody solutions were prepared fresh in PBS, 100 µl added to each section and allowed to bind overnight at 4°C (primary) or 1 h at room temperature (secondary). Details of antibody dilutions and detection method for all targets and detection conjugates can be found in Table S1.

After secondary antibody incubation, slides were washed 3 times for 5 min each in PBS. For biotinylated secondary antibodies, this was followed by a 10 min incubation with extravidin peroxidase (Sigma, E2886), 1:100 in PBS, then rinsed in PBS, before being developed using DAB (Abcam, ab64238) for 10 min, prepared according to manufacturer’s instructions. Slides were washed 3 times for 5 min each in PBS and counterstained with nuclear fast red (Molecular Probes, R37106) for 5 min in the fume hood, Haematoxylin for 3 min or Eosin for 30 seconds, then rinsed in PBS and dehydrated in ethanol, 95 % for 1 min followed by three times in 100 % for 10 dips in each and immersed twice in Histoclear for 3 min each. Dehydration steps were not carried out for PCNA stain. Slides were left to dry for a few min and mounted with quick-drying mounting media, or glycerol jelly mounting media secured with nail polish. Typically, the stains above were carried for each of n = 3 mice per experimental group, x 4 tissue sections per mouse, x 4 random images per tissue section, typically n= 48 datapoints per experimental group. Any deviations from this are made clear in figure legends.

### 4T1GFP+Luc Cell Retrieval from Tumour-Bearing Mice

Tissues (brain, lung, kidney, liver) were collected following perfusion with 2 mM EDTA in PBS, into ice-cold DMEM (Gibco, 41965-039) on days 3 and 7 post-implantation. Lungs, kidneys and Livers were digested in 3 mL 0.2% Collagenase A (Roche, 11088793001) in PBS for 30 to 45 min at 37 °C. Cell suspensions were filtered through a 70 µm mesh and centrifuged at 260 x g for 5 min. Pellets with high RBC content were treated with 2 mL ACK Lysis Buffer (Gibco, A1049201) for 2 min, neutralised with 8 mL PBS, and re-centrifuged. Brains were minced and digested in 2-3 mL 0.2% Collagenase A containing 10 µg/mL DNase I (Roche, 10104159001) for 30 min at 37 °C, then centrifuged at 280 x g. Pellets were resuspended in 10 mL filter-sterilised 20% BSA-DMEM (Sigma, A7906) to remove myelin, mixed gently (20 passes), and centrifuged at 1000 x g for 10 min to remove the myelin layer. The supernatant was discarded, pellets resuspended in 2 mL DMEM, filtered through a 70 µm strainer, and re-centrifuged at 280 x g. Cell pellets were resuspended and plated in 6-well plates (one well/mouse) in RPMI (Gibco, 21875-034) + 10 % FBS (Gibco, 10270-106) + 60 μM 6-Thioguanine (ThermoFisher, B21280.06). 6-TG containing medium was replaced the next day and selection maintained for 2 more days to eliminate non-tumour cells.

Attached cells were rinsed with PBS and then either stained with crystal violet in 70% ethanol for 30 min, washed and air dried for imaging, or fixed with 70% ethanol for 8 min and stained with nuclear fast red for 10 min, washed twice with diH O, and air-dried before imaging. Plates were imaged using an EVOS Fl Auto 2 scanner at 10 x. Colonies were counted manually and those comprising ≥ 5 cells were classified as “large”. Parallel experiments were performed with tissues from tumour-free mice as controls.

### Imaging and Analysis

H&E and IHC sections were imaged in brightfield using an Olympus BF microscope or Leica DM5500. Immunofluorescence was acquired on a Leica DM5500 epifluorescence microscope.

IHC signals (SNAIL, Vimentin, Arg1, NOS2, CD3, MAC2) were quantified using QuPath(Bankhead et al., 2017) (v0.2.3). Test sections were compared to negative controls on the same slide using a 1 x 2 grid view. Stain vectors were set using annotations of signal, counterstain, and background. A threshold was applied (Gaussian prefilter, sigma = 1, initial threshold = 0.35) and adjusted to detect signal above threshold.

Magnification used was adjusted to each target, and is clearly described in figure legends. For PCNA staining, PCNA-positive cells were counted using QuPath’s positive cell detection plugin. Due to faint nuclear staining, proliferation was also estimated in FIJI as the ratio of PCNA-positive area to total tumour area (measured at 4 x).

Optical Density and necrosis were quantified using FIJI (Schindelin et al., 2012), where images were converted to 8-bit greyscale, then tumour and background means were measured and background subtracted from tumour mean. Necrotic areas (white, friable, granular) were measured using the free-hand tool at 4 x. Necrosis ratio = [Total necrotic area / Total tumour area].

Tumour cell colony images were imported into FIJI and colonies were counted blindly and manually using the multi-point tool. Proliferation potential was calculated as the ratio of large colonies (≥ 5 cells) to total colonies.

## Data Analysis

Data were analysed using Microsoft Excel 2010, GraphPad Prism (v9.0.2 or v10.2.3), and QuPath (v0.2.3). Statistical comparisons between experimental groups were performed using nested one-way ANOVA followed by Tukey’s multiple comparisons test, accounting for multiple measurements obtained from individual biological replicates (mouse). The mouse was considered the independent experimental unit, with tissue sections and image fields treated as subsamples rather than independent biological replicates. Comparisons between two independent time-point cohorts within the same implantation-density group were performed using nested unpaired t tests.

The relationship between HIF-1α and PDGFRβ staining was assessed using Pearson’s correlation coefficient, calculated from paired mean values obtained from individual tumours (n = 9 biological replicates per time point, comprising 3 mice from each implantation density group). Correlations were analysed separately for tumours collected at 3 d and 21 d post implantation, using two-tailed significance testing.

The specific statistical analyses applied to each parameter are indicated in the corresponding figure legends.

## Results

To assess how different tumour induction protocols for TNBC affect the tumour microenvironment and the biology of these tumours, we induced orthotopic mammary tumours using 4T1^GFP+Luc^ cells (Wright et al., 2016) into the 4^th^ (inguinal) mammary fat pad of 8-week-old female Balb/c mice, by implanting 1 × 10^4^, 5 × 10^4^, or 2.5 × 10^5^ cells. Mouse age was selected to represent the typical onset of human TNBC in human patients, and the cell numbers used are at the lower end of the reported range reported in the literature, where tumours are mostly generated with ≥ 5 × 10^5^ cells. Primary tumours were excised after 3, 21 and 35 d after implantation. FFPE sections were stained with H&E, and IHC was used to characterise multiple parameters of the TME at different stages of tumour development.

### Number of cells implanted does not impact tumour volume at early stages, but affects tumour density, necrosis and hypoxia

The tumour progression was surprisingly consistent across experimental groups, which were all palpable by 21 days after implantation. Tumour volume (Fig. 1A) and tumour section area (Fig. S1A, B) are not affected by number of cells implanted until advanced stages. Tumours generated with the highest number of cells (2.5 × 10^5^) became significantly larger than the two other groups only from day 31 onwards. The number of cells injected did not significantly affect the host weight over time (Fig. S1C) or tumour burden (tumour-to-body weight ratio, Fig. S1D). We queried why tumours would not follow a linear growth progression (proportional to the number of cells implanted) and postulated that distinct parameters within the TME would underpin their growth kinetics. Using H&E-stained sections imaged at low magnification (Fig. 1B), the average optical density was measured, informing that at 3 d and 21 d after implantation, tumours initiated with 2.5 × 10^5^ cells were markedly and significantly denser than the two tumour groups generated with fewer cells; this significance is no longer seen in tumours excised after 35 d (Fig. 1C) where all tumour groups show comparable optical densities. Using FIJI we were able to evaluate and quantify the necrotic area fraction in the same tissue sections. When these data are shown as a function of cell density for each time point (Fig. S2A), the necrotic areas are not significantly different between groups in tumours excised at 3 d, but tumours generated with 5 × 10^4^ and 2.5 × 10^5^ cells show higher necrotic area fractions than those from the 1 × 10^4^ cells group after 21 d, once tumours are established, although this difference only reaches statistical significance at more advances stages (35 d). When displayed as a function of time for each experimental group (Fig. 1D), it becomes apparent that the baseline necrotic area fraction in tumours generated with 1 × 10^4^ and 5 × 10^4^ cells does not significantly increase over time (Fig. 1D, left and middle panels), whereas necrosis is significantly higher in tumours at advanced stage (35d) when generated with 2.5 × 10^5^ cells (Fig. 1D, right panel). Taken together, these data show that while both 5 × 10^4^ and 2.5 × 10^5^ tumours are more necrotic than those generated with fewer cells, only the ones in the densest cohort have progressively increased necrosis as the tumour develops. Interestingly, for tumours generated using 5 × 10^4^ cells, the observed increase in necrosis is not mirrored by a similar increase in optical density.

**Fig. 1.**
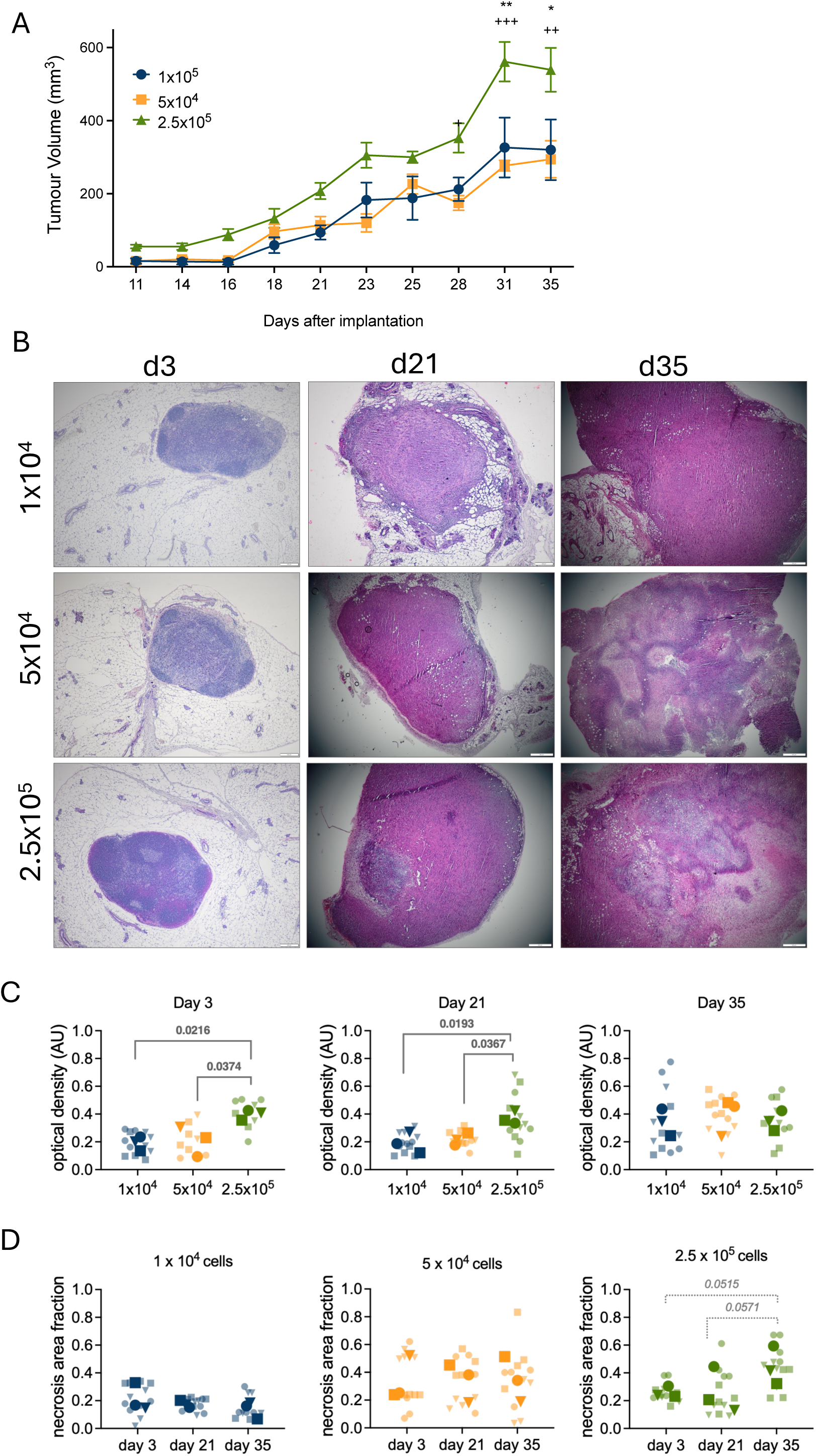
Effect of cell number on tumour growth, density and necrosis. Primary mammary tumours generated with different numbers of 4T1 cells (1 × 10^4^, 5 × 10^4^, or 2.5 × 10^5^) (A) Tumour volumes were recorded for the duration of the experiment and are plotted as average ± StDev, statistical significance between groups was assessed by one-way ANOVA at each time point; * refers to 2.5 × 10^5^ (green) vs 5 × 10^4^ (yellow) and ^+^ to 2.5 × 10^5^ (green) vs 1 × 10^4^ (blue); (B) Representative images of H&E sections from mammary tumours excised 3, 21 and 35 d after implantation; Scale bar = 500 μm (C) Optical density of tumour sections was calculated using FIJI. n= 3 tumours per experimental group, x 4 sections per tumour; 12 datapoints total per experimental group (faded symbols) and average for each tumour (solid symbols) are shown; (D). Average area of necrosis in primary tumours was calculated using FIJI. As in C, faded symbols represent images from the same section of the tumour (average shown in solid symbols); data plotted for each experimental group over time (see Fig.S2 for comparisons between groups within each time point). For C and D, statistical significance was assessed by *nested* one-way ANOVA p values shown in comparisons that were significant or approached threshold for significance (*italicised)*.

The large necrotic cores seen in advanced stage tumours (35 d) caused challenges in sectioning and staining, and carried the added caveat of quantifying signals in non-viable cells. Thus, to avoid confounding the differential characterisation of the individual tumour cohorts, the subsequent parameters were assessed exclusively in 3 d and 21 d tumours, representing incipient and established tumour stages, and which are key for tumour cell evolution and systemic priming.

### Tumour proliferation rate is inversely proportional to the number of cells used for implantation

To assess cell proliferation in tumours initiated from different cell numbers, we stained tumour sections for PCNA (Sarraf et al., 1991). Proliferating cells were expected primarily in the periphery of the tumour (Gallaher et al., 2019), but were instead often found in defined clusters close to the edge of the tumour (Fig. 2A). In tumours excised 3 d post implantation, the number of PCNA positive cells is higher in the tumours generated with the lowest number of cells (1 × 10^4^) compared to 5 × 10^4^ and 2.5 × 10^5^ (Fig. 2B). Due to faint nuclear staining in these sections, it was not possible to accurately quantify PCNA-negative cells and to normalise to the total number of cells per image. Therefore, we also quantified the percent proliferation area (Fig. 2C, area of PCNA-staining/total area of section in field of view × 100). While the number of proliferating (PCNA-positive) cells between groups (Fig. 2D) is not statistically different, the proliferation area remains highest in the tumours generated with 1 × 10^4^ cells (Fig. 2E). These observations were further validated with an additional proliferation marker, Ki67 (Davey et al., 2021), used in consecutive sections of the same tumours (data not shown). Together, these readouts show that tumours generated with the fewest cells (1 × 10 ) exhibit greater proliferation, with statistically significant differences in PCNA-positive area compared with the other implantation groups at both early (3 d) and established (21 d) stages, determining that tumour cell proliferation rate progresses inversely to the number of cells used for implantation.

**Fig. 2.**
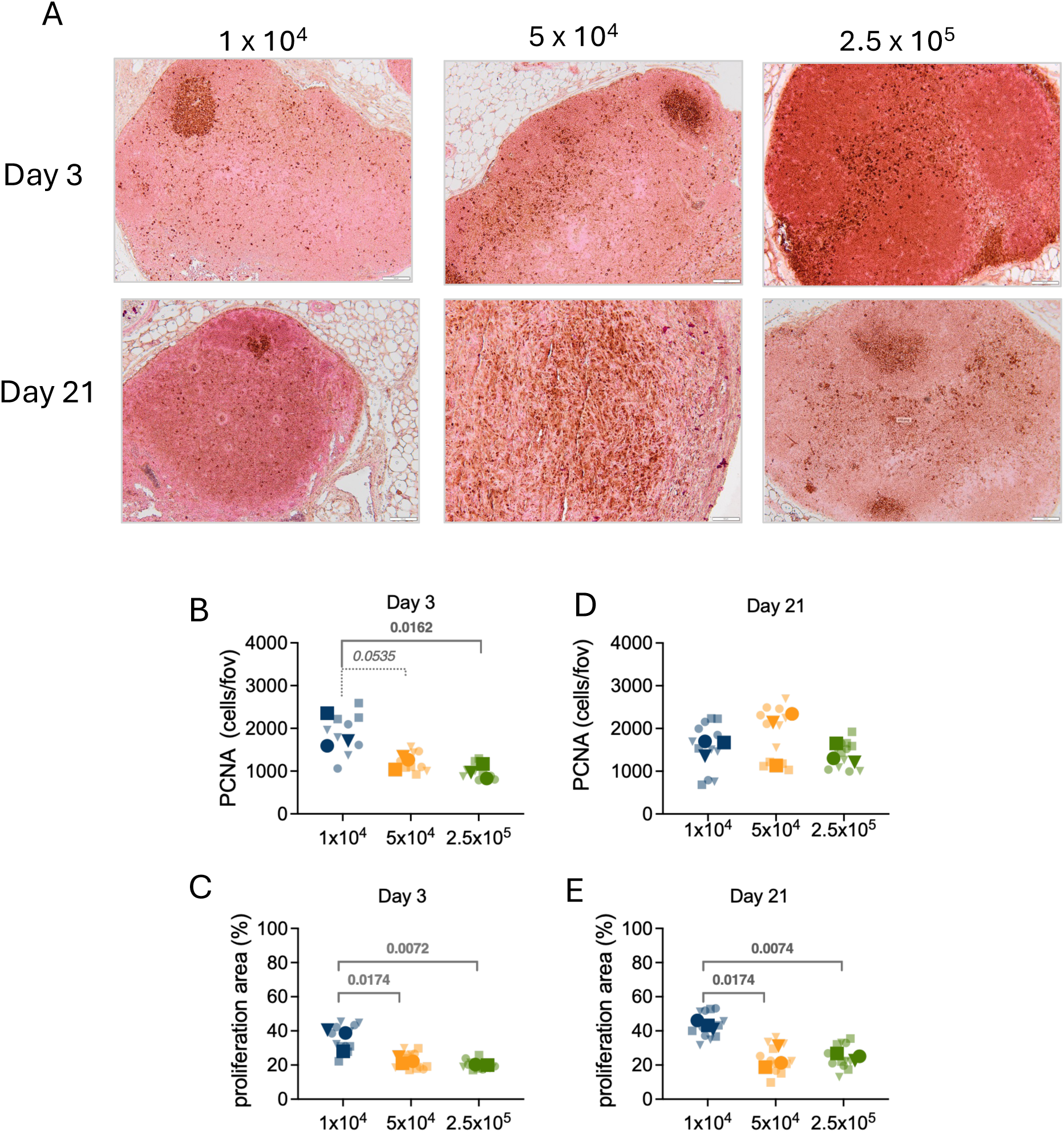
Proliferation rate using PCNA staining of primary tumours. Mammary tumours generated with different number of 4T1 cells (1 × 10^4^, 5 × 10^4^, or 2.5 × 10^5^) were excised 3 and 21 d post implantation into mammary glands. (A) Representative images of IHC primary tumours sections probed with anti-PCNA antibody. Scale bar = 100 μm. Total number of cells stained with PCNA in 3 d, shown as number of cells per field of view (fov) (B) or shown as a percentage of area of PCNA signal/total section area C); Panels (D) and (E) represent the same data as in B and C but for 21d tumours. n= 3 tumours per experimental group, x 4 sections per tumour, n = 12 datapoints per experimental group/timepoint. For panels B-E, shaded symbols indicate individual values (for each section), solid symbols of same shape represent average for that tumour; Statistical significance was assessed by *nested* one-way ANOVA p values shown for comparisons that reached or approached threshold for significance (*italicised)*.

Highly proliferative tumours can cause necrosis as they become hypoxic and often deficiently vascularised (Mus et al., 2015). We quantified HIF-1α (Fig. 3A, S3A) and CD31 (Fig. 3B and Fig S3B), as markers of hypoxia and endothelial cells, respectively, in sections from the different tumour groups over time. Tumours generated with the lowest number of cells (1 × 10^4^) were significantly less hypoxic than those generated with 5 × 10^4^, and 2.5 × 10^5^ cells, and this is statistically significant at both 3 d and 21 d. These data indicate that tumour hypoxia does not necessarily result in necrosis; tumours generated with 5 × 10^4^ and 2.5 × 10^5^ have significantly higher proportion of cells with stable HIF-1α (Fig. 3A), however only the latter has significantly larger necrotic areas compared to the remaining two groups (Fig. 1C). This suggests that the threshold at which the response to hypoxia is sufficient and effective in preserving cell viability is crossed at a specific cell density, which in this model is between 5 × 10^4^ and 2.5 × 10^5^ cells.

**Fig. 3.**
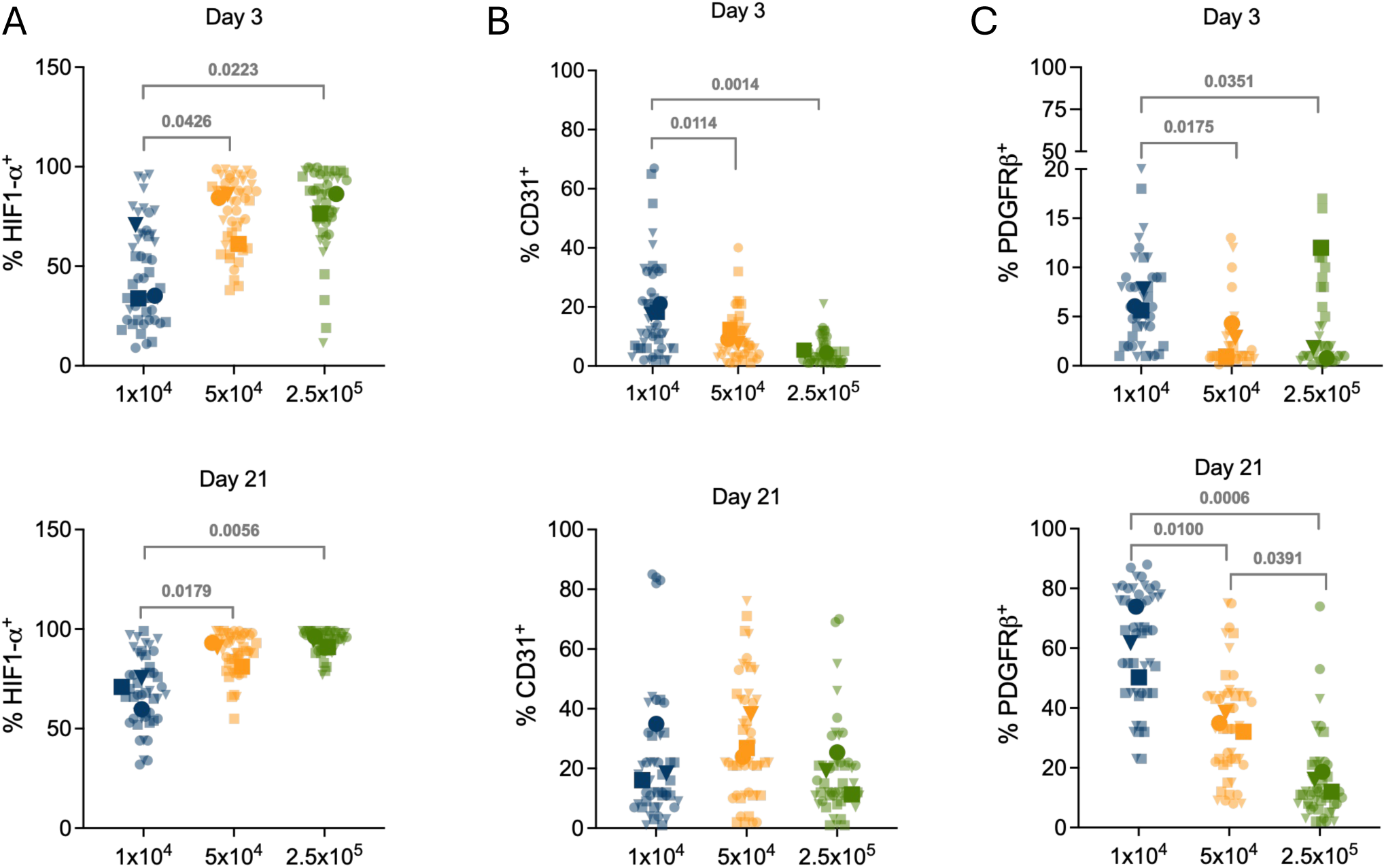
Hypoxia, vascular density and vessel integrity in different tumour protocols. Mammary tumours generated with different number of 4T1 cells (1 × 10^4^, 5 × 10^4^, or 2.5 × 10^5^) were excised 3 and 21 d post implantation. Cells were stained for (A) HIF-1α as a hypoxia marker (B) CD31 for endothelial cells and (C) PDGFRβ as a marker for tumour pericytes. QuPath was used to calculate either percent area of signal (A) or percent number of cells normalised to nuclear stain (B and C); n= 3 mice per experimental group, x 4 sections tumour, x 4 images per tissue section, n = 48 datapoints total per experimental group; faded symbols represent individual values for each image, solid symbols of same shape show the average for each tumour. Statistical significance was assessed by *nested* one-way ANOVA p values shown for comparisons that reached or approached threshold for significance (*italicised)*.

Interestingly, the proportion of CD31+ cells (Fig. 3B) is only significantly higher in the lowest cell density group, and at 3 d post implantation (Fig. 3B, upper panel). In 21 d tumours this parameter is no longer different between experimental groups (Fig. 3B, lower panel), indicating that high levels of HIF-1α and/or hypoxia seen in these same tumours are not a result of lower vascular density, since nearly all cells display some level of HIF-1α accumulation.

Pericytes are stabilisers of vascular integrity, and loss of pericyte coverage is one well-characterised feature of tumour vascular dysfunction (Craggs et al., 2015; Yao, 2022). We stained consecutive sections from those stained for CD31, with PDGFRβ (Hosaka et al., 2016), a commonly used marker for tumour pericytes, typically staining those in association with endothelial cells, and thus appropriate to infer pericyte coverage and vessel integrity.

In 3 d tumours, very few PDGFRβ+ cells are detected (Fig. 3C, upper panel, *note split y-axis*, and Fig. S3C), in keeping with the incipient, immature microvasculature expected in early tumour development. We further assessed the relationship between HIF-1α and PDGFRβ staining using paired measurements from individual tumours (Fig. S3D). At 3 d post implantation, a negative but not statistically significant correlation was observed (Pearson r = −0.5498, p = 0.1251, n = 9 tumours). In the 21 d cohort a stronger inverse correlation was observed, which reached statistical significance (Pearson r = -0.8853, p = 0.0015). These observations suggest that differences in HIF-1α and PDGFRβ staining may reflect differences in vascular maturity, although vascular integrity and perfusion were not directly assessed.

This result aligns with the hypoxia reading in Fig. 3A, as reduced pericyte coverage is associated with unstable and dysfunctional vasculature (Nagy et al., 2009). Combined, these data indicate that the increased HIF-1α signal observed in the tumours generated with higher cell numbers (Fig. 3A) is more likely a reflection of differences in vascular integrity and perfusion (Fig. 3C) rather than reduced endothelial cell abundance (Fig. 3B).

### Presence of immune cell infiltration increases with cancer cell number

Necrosis and hypoxia are both triggers of inflammation (Bartels et al., 2013; Finger & Giaccia, 2010), and we investigated if recruitment of macrophages to the primary tumour was influenced by the number of cells used to generate it. We used MAC2 (Galectin-3) as a marker for tumour associated macrophages which typically display an M2-phenotype (Sciacchitano et al., 2018; Zhu et al., 2021). In this analysis, the signal at the margins (periphery) and the centre (core) of each tumour section were qualified (Fig. 4A) to interrogate how the presence of these cells, as well as their ability to penetrate the tumour, was influenced by the inducing protocol over time. Peripheral MAC2+ cells increase with tumour progression in all experimental groups (Fig. 4B), and more macrophages are seen in tumours generated with higher numbers of cells (Fig. S4A). Fewer MAC2+ cells are found in the tumour core (Fig. 4C), but the trend mirrors that seen at the margins and an increase in macrophage numbers as tumours grow; very few MAC2+ cells are detected in the tumours generated with fewest cells (1 × 10^4^) (Fig.4C, left panel, note split *y*-axis). These same data plotted as a function of cell number (Fig. S4A) show that increased tumour cell density is associated with MAC2+ macrophage recruitment, particularly at the tumour periphery. Quantification of pan-macrophage marker CD68 (Song et al., 2023) in whole tumour sections (Fig. S4B, left panel) confirms the trend seen for MAC2, suggesting a link between the increased tumour size, cell density and necrosis with the presence of myeloid-derived cells. We complemented this by assessing the presence and density of T lymphocytes in the tumours using CD3 as a marker (Lee et al., 2017). In early tumours the presence of T-cells is not significantly affected (Fig. S4C, right panel), but by 21 days, tumours generated with 5 × 10^4^ and 2.5 × 10^5^ cells show significantly higher number of CD3+ cells compared to the 1 × 10^4^ group.

**Fig. 4.**
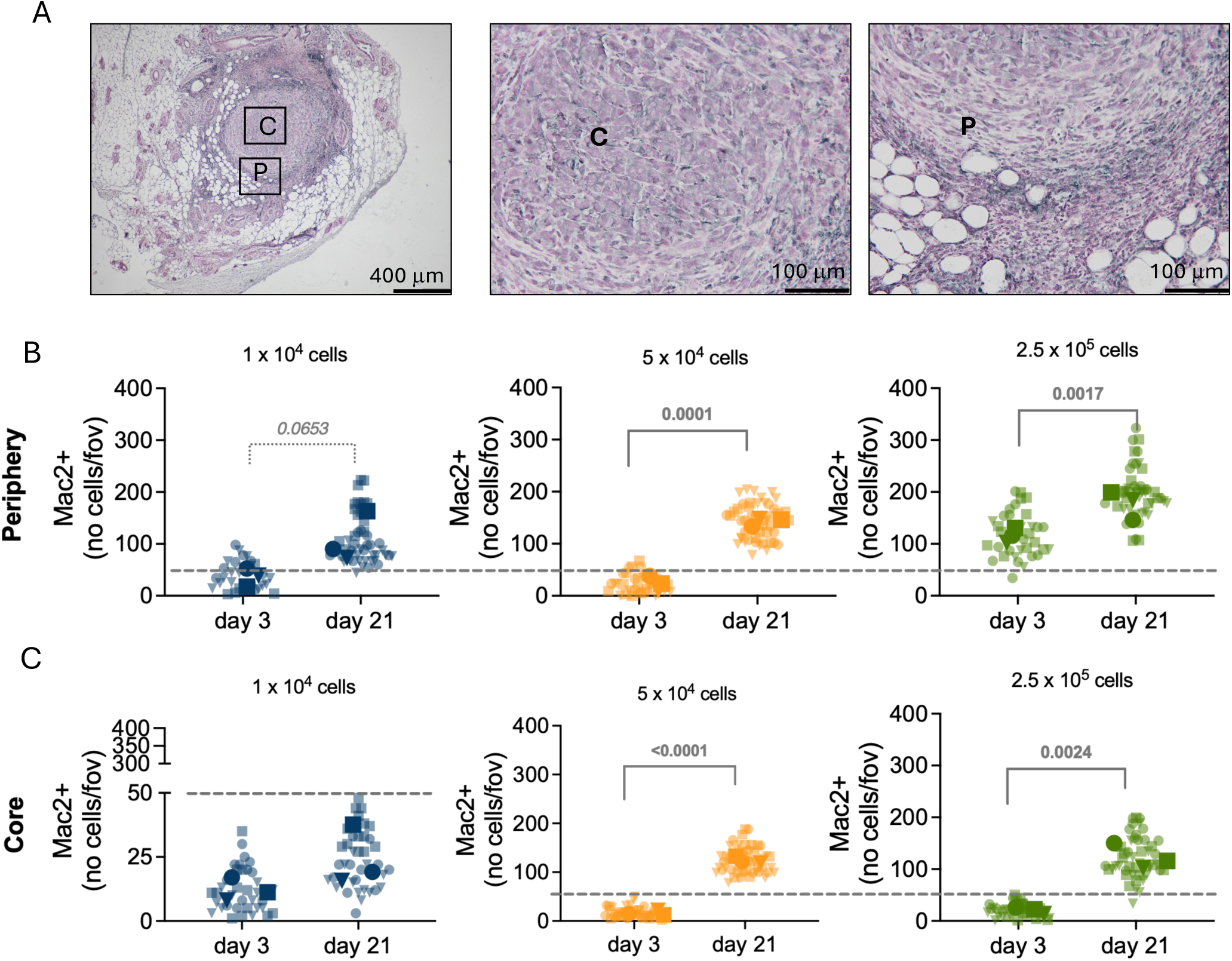
Number of MAC2+ cells increase with tumour progression. (A) Representative qualification of core and peripheral tumour areas used to quantify the presence of MAC2+ cells. The number of cells per field of view (fov) of peripheral (B) and core (C) macrophages were counted in early (3 d) and established (21 d) tumours in each experimental group – 1 × 10^4^ (blue), 5 × 10^4^ (yellow), and 2.5 × 10^5^ cells (green); each symbol shape represents one biological replicate (mouse) with faded shapes depicting each image readout, and corresponding symbols identical symbols in solid colour indicating the average value for each tumour; n = 3 tumours, ≥ 4 sections stained per tumour, ≥ 4 random images obtained for quantification (minimum 48 datapoints per tumour); Statistical significance between 3 d and 21 d tumours within each implantation-density group was assessed using nested unpaired t-tests, accounting for multiple measurements obtained from individual mice; p values shown for comparisons that reached or approached (*italicised)* threshold for significance.

### Influence of tumour induction protocol, tumour microenvironment and dissemination potential

One of the main challenges in modelling breast cancer progression has been studying its organotypic dissemination. Organ-specific tropism can be created by repeated metastatic cell selection and retrieval and reimplantation, increasingly enriching for organ-specific colonisers (Nakayama et al., 2021) or by direct infusion into specific vascular beds: intracardiac for brain or skeletal metastasis (Berce, 2025;

Terrell-Hall et al., 2017; Wright et al., 2016), carotid artery (brain metastasis and limiting risk of extracranial tumours) (Lim et al., 2022), tail vein frequently used for lung colonization (Branco-Price et al., 2012; Shrestha et al., 2019), or hepatic portal vein, for modelling liver metastasis (Goddard et al., 2016). These models of experimental metastasis, however, sidestep the initial steps of the spontaneous dissemination cascade (Labelle & Hynes, 2012), primarily that of tumour-priming of distant organs. We briefly looked at markers of EMT SNAIL (Kim et al., 2016) and Vimentin (Yamashita et al., 2013) in tumour sections from each of the experimental groups (Fig. S4E). While some differences appear statistically significant, there was no obvious link to the number of cells used to generate the tumours, and results proved inconclusive to investigate likelihood of association between the induction protocol, EMT and dissemination. This experiment was repeated aiming to focus on the earlier stages of tumour progression, as TNBC is frequently disseminated at the time of diagnosis. As before, 1 × 10^4^, 5 × 10^4^ and 2.5 × 10^5^ 4T1 cells were implanted into a single mammary gland, and the experiment was terminated 3 days and 7 days later. Metastatic (brain, lung and liver) and non-metastatic (kidney) visceral organs were collected, digested into a single cell suspension, and tumour cells selected in medium containing 6-thiguanine (Pulaski & Ostrand-Rosenberg, 2001). Colonies were fixed, stained and counted, and classified between small (up to 4 cells) and large (5 cells or more) to gauge their proliferative potential and not just their survival post isolation (Fig. S5A). The relative number of total colonies isolated from each organ 3d post implantation in each experimental group is shown in Fig. S5B. While our observations are not statistically significant, they proved informative for several parameters. Firstly, that tumour cells disseminate as early as 3 d from the primary tumour to multiple metastatic and non-metastatic (kidney) organs. This was unexpected, especially because we do not typically observe overt metastasis within this experimental timeframe, and pulmonary metastases remain the most common, albeit seen in much more advanced tumour stages. At 3 d post implantation, a higher proportion of colonies recovered from brain and lung tissues originated from tumours generated with the fewest cells (1 × 10 cells), although these differences were not statistically significant. These observations reflect the recovery and ex vivo expansion of viable tumour cells rather than a direct measure of their dissemination efficiency or organ-specific colonisation.

For isolations performed 7 d post implantation (Fig. S5C), most colonies obtained from brain still originated from the 1 × 10^4^ tumour group. The colonies obtained from lung tissue were equally distributed across experimental groups. Very few colonies were obtained from the 5 × 10 group, an intriguing observation that warrants further investigation in larger experimental cohorts.

To investigate if the tumour cells isolated from different organs carried differential ability to proliferate, the number of large colonies (≥ 5 cells) was presented as a fraction of the total. In most tissues, tumour cells isolated from the least dense tumours showed a tendency to form larger colonies (Fig. S5D), both at 3 d or 7 d post implantation, although these differences were not statistically significant. Exceptionally, large colonies generated from lung tissue and 7 d post implantation are similarly distributed across experimental groups, whereas very few large colonies are found in the livers of mice bearing tumours from 5 × 10^4^ cells group. This experiment remains exploratory and should not, as presented, be interpreted as evidence of organotropism or differences in dissemination efficiency. The number and size of colonies recovered may reflect several processes, beyond the presence of tumour cells in or associated with the harvested organs, such as their survival during tissue processing, recovery under 6-thioguanine selection, and subsequent proliferation *ex vivo*. These observations do not establish whether the recovered cells would successfully form metastases *in vivo*. Nevertheless, they provide insight into opportunities to use pre-clinical models and protocols to investigate early dissemination and growth potential in different environments.

## Discussion

### Tumour cell number is a primary determinant of tumour microenvironmental state

Orthotopic models of breast cancer are essential to investigate mechanisms of progression and to test strategies to treat the disease, as they generate a single tumour within the appropriate tissue context of the mammary gland. However, studies report a vast range of methods to create mammary tumours by implantation, even within the 4T1 model of TNBC. Thus, whether experimental outcomes primarily reflect the tumour biology or the method used to study it remains a key unresolved question. For the collective understanding of breast cancer progression, it is critical that experimental observations are interpreted within the context of how tumours are generated, since, as shown here, this will have fundamental implications in the tumour microenvironment.

In this study, using the established 4T1^GFPLuc^ model (Pulaski & Ostrand-Rosenberg, 2001) the number of tumour cells introduced into the mammary gland was the only variable considered, while preserving remaining parameters constant (e.g., age, injection volume, carrier/matrix). We show that tumours induced with distinct cell numbers become palpable within the same timeframe and remain largely indistinguishable by *in vivo* measurements and FFPE section area until advanced stages. Only tumours generated with the highest number of cells become significantly larger at later time points. Despite these similarities in gross growth kinetics, tumours differed markedly in density, hypoxia-associated responses, necrosis, proliferation, vascular-associated markers, and immune infiltration. These findings indicate that tumour induction protocol dictates tumour microenvironmental state more profoundly than tumour size or time since implantation. This understanding is an important consideration for relevant and useful experimental design.

### Vascular markers suggest differences in vessel integrity associated with hypoxia

TNBC tumours are typically more densely vascularised than those in other breast cancer subtypes (D’Andrea et al., 2021; Mirando et al., 2020). While vascularization, as measured by CD31 staining, was more efficient in low-density tumours at early stages, microvascular density is not distinguishable between experimental groups once tumours become established (21 d). Despite comparable apparent endothelial cell staining, HIF-1α expression was significantly lower in tumours generated with fewer cells. The increased HIF-1α signal was associated with reduced PDGFRβ staining, potentially reflecting differences in vessel maturity (D’Andrea et al., 2021). Combined, these observations suggest that hypoxia in tumours induced with higher cell density may reflect impaired vessel integrity and perfusion rather than reduced vascularization. The implications of this are relevant, as differences in vascular function, including early access to functional vasculature in low-density tumours has the potential to influence tumour cell behaviour and metastatic properties, including dissemination efficiency and viability at distant sites. Hypoxia is central to modulation of the tumour microenvironment, and implicated in metabolic adaptations: the associated increase in glycolytic activity contributes to EMT (Hapke & Haake, 2020), treatment resistance and immunosuppression (Zhou & Zhou, 2025). Primary tumour hypoxia is also known to influence metastatic patterns (Rankin & Giaccia, 2016) and tumour cell fate at metastatic sites (Fluegen et al., 2017). Our findings underscore how subtle differences in induction protocols can fundamentally alter tumour physiology and disease progression from the earliest stages of tumour onset.

### Immune infiltration and inflammatory features scale with tumour induction density

Necrosis and hypoxia are well-established triggers of inflammation(Bartels et al., 2013) and in breast cancer necrosis correlates with advanced disease and macrophage infiltration(Leek et al., 1999). Consistent with this, tumours generated with higher numbers of cells developed substantially larger necrotic areas and exhibited significantly greater macrophage recruitment (MAC2 staining), more densely at the tumour periphery but also within the tumour core. In contrast, tumours generated with fewer cells remained largely proliferative, showed minimal necrosis, and expectedly exhibited limited macrophage infiltration.

These differences in immune cell recruitment were supported by additional pan-macrophage (CD68) and T-cell (CD3) markers, firming the relationship between tumour cell density and factors affecting inflammatory and immune landscape of the primary tumour. These parameters are intrinsically dependent on a dynamic vascular profile, and influence not only the tumour cell, but the whole local and systemic ecosystem, in response to secreted factors; The type and quantity of cytokine and chemokine signals generated are expected to reciprocally contribute to vascular and extracellular matrix patterns, metabolic demands and metastatic priming (D’Andrea et al., 2021; Guo et al., 2024; Zhang et al., 2025). Furthermore, the kind and extent of immune infiltration critically influences therapeutic response (Zhang et al., 2023; Zhang et al., 2025; Zhu et al., 2021), underscoring the implications of experimental design in immunological readouts, and potential for hindering cross-study comparisons, even when using nominally identical tumour models.

### Early tumour cell dissemination is observed across implantation protocols

One of the major challenges in modelling breast cancer progression is capturing the earliest steps of spontaneous dissemination, which are critically influenced by signals originating in the primary tumour and by host responses to tumour initiation(Labelle & Hynes, 2012). TNBC is the most metastatic of breast cancers(Anders & Carey, 2009; Dent et al., 2007), and this is the primary cause of mortality in these patients. While advanced tumour stages are typically associated with more invasive phenotypes, established breast cancer metastases have been shown to originate from early disseminated tumour cells instead of from mature tumours (Hosseini et al., 2016). Metastatic dissemination can occur before the primary tumour is detectable, and indeed TNBC patients often have disseminated disease at the time of diagnosis and/or very early relapse, primarily to visceral organs(Anders & Carey, 2009). Therefore, we investigated we investigated whether viable tumour cells could be recovered from distant organs at the earliest stages post implantation. Tumour cells were consistently retrieved from multiple organs as early as 3 d post implantation across all tumour induction protocols, indicating that early dissemination is a common feature of orthotopic 4T1 tumours. Qualitative differences in the recovery and ex vivo colony formation of tumour cells were observed across implantation conditions, providing an opportunity to explore how tumour induction protocols may influence early dissemination-associated processes. However, these observations cannot distinguish differences in tumour cell dissemination from differences in survival, recovery or subsequent proliferation following isolation, and indeed if allowed to persist in the metastatic organ. Cells originating from low-density tumours (1 × 10 ) were more frequently recovered as colonies from brain and lung tissues at early time points and showed a tendency to form larger colonies upon isolation, although these differences were not statistically significant. The recovery of tumour cells from distant organs is influenced by several processes that cannot be distinguished using this experimental approach. Differences in colony number may reflect variations in the number of cells reaching the organs, their retention or survival within the tissue, or their ability to withstand tissue processing and subsequent selection in culture. Similarly, the formation of larger colonies may indicate differences in proliferative capacity under *ex vivo* conditions, but does not necessarily predict successful metastatic colonisation *in vivo*. Thus, while these observations suggest potentially interesting differences between tumour induction protocols, further experiments would be required to establish the mechanisms involved.

The properties of the early tumour microenvironment (hypoxia, necrosis, macrophage infiltration, vascular integrity) are consequential for tumour cell phenotypes, but also secretory and signalling profiles that will shape host-mediated responses to disseminating cells (Finger & Giaccia, 2010). It is unclear then what underpins early dissemination: the priming of the tumour cell or that of the distant, premetastatic organ, in each of these protocols. To investigate this, assessment of tumour-derived factors, such as by whole tumour digestion and secretome, characterization of plasma cytokine profile and validation of corresponding remodelling at pre- or early metastatic organs is necessary, in much larger and adequately powered experimental cohorts, which befalls beyond the scope of this study.

We briefly assessed SNAIL(Kim et al., 2016; Lundgren et al., 2007) and Vimentin(Grasset et al., 2022; Kim et al., 2016; Yamashita et al., 2013) in primary tumours, to reflect EMT-associated markers reported as critical for metastatic success in breast cancer. The results do not show a linear density-dependent trend precluding conclusive interpretations. This aligns with previously reported data (Grasset et al., 2022) showing that, in TNBC, most cells within primary and metastatic tumours are in a dynamic and frequently hybrid endothelial/mesenchymal state; therefore, single stains for either state are insufficient to define the tumour cell invasive phenotype, or predict progression.

Our data, however, indicate that a complex crosstalk is at play between tumour cell density, microenvironmental stress, and host responses. One possibility raised by these observations is that differences in tumour induction density may influence early host responses, including innate immune surveillance of circulating tumour cells. However, alternative explanations include differences in tumour cell survival, retention within distant tissues, or subsequent expansion under *ex vivo* selection conditions. The present experiments cannot distinguish between these possibilities. While not directly tested in this study, the potential contribution of host responses is consistent with the observed differences in necrosis and inflammatory infiltration between tumour induction protocols, and warrants further investigation. This information can be leveraged and used as a tool to design experiments and create models most appropriate to investigate processes governing early dissemination events.

Furthermore, metabolic priming of primary tumour cells, as well as their plasticity at distant sites is fundamental for metastatic progression and cell fate post dissemination (Fluegen et al., 2017; Roshanzamir et al., 2022). Together, these findings demonstrate that tumour induction protocols have profound consequences for tumour microenvironmental development, with implications for early tumour cell dissemination and behaviour. While the mechanisms underlying the observed differences in colony recovery remain to be established, these observations reinforce the importance of tumour induction protocols in shaping experimental outcomes and their biological interpretation.

### Implications for model selection, reproducibility, and translational relevance

Models of spontaneous metastasis are often designed to prioritise metastatic readouts, frequently requiring primary tumour resection(Tai et al., 2014) or prolonged experimental timelines, leading to either additional procedures or heavy tumour burdens, with negative implications for experimental animal welfare (Paschall & Liu, 2016). However, the composition of the primary tumour will shape the signals distributed systemically, more strongly by tumour cell density than by tumour size or time since implantation, which cannot be assumed to represent tumour stage. Diffusion and efficacy of therapeutics is affected by the biological (Galardi et al., 2025; Lasso et al., 2019; Zhou & Zhou, 2025) and physical properties of the microenvironment, with, for example, high-density tissues hindering diffusion (Kim et al., 2017; Sabit et al., 2025). This leads to selective killing of some tumour cells, leaving others intact and able to evolve, disseminate and colonise. Inevitably, variability in tumour induction protocols will contribute to inconsistent therapeutic responses across preclinical studies and, ultimately, compromise success in trials and progression to the clinic. Intentional model design, aligned with specific biological questions and disease stages, is therefore indispensable to improve reproducibility and translational relevance. Understanding the specific properties generated under each protocol will allow refinements to match specific clinical states and stages. Importantly, it is important to note that pre-clinical murine cancer models are inherently biased towards lung colonisation, which results in extreme morbidity before metastasis in other organs can develop. Circumventing this requires resection, organotropic selection and/or manipulation of tumour cells in advance of implantation. Our results demonstrate that tumour cells can be recovered from multiple distant organs at early time points, while highlighting the potential importance of primary tumour microenvironmental cues in shaping their subsequent behaviour. This can be leveraged to investigate circumstances in which cells are more or less likely to survive and thrive in organs that they are able to reach. For example, by modelling preexisting conditions, or the effect of environmental and therapeutic cues.

### Study limitations and opportunities for future investigation

This study was designed to isolate the impact of tumour cell number at implantation, and several limitations should be acknowledged. First, while early dissemination was reproducibly observed, these experiments were exploratory and not powered to quantitatively assess dissemination efficiency or immune-mediated clearance. Future studies incorporating larger cohorts, immune cell profiling, and functional depletion approaches will be required to directly test whether tumour cell density modulates early immune surveillance.

Second, assessment of EMT was limited to immunohistochemical detection of SNAIL and Vimentin, which proved insufficient to capture dynamic, hybrid, or transient invasive states. More informative approaches would include flow cytometric analysis of GFP-labelled tumour cells, transcriptional profiling at early time points, and functional invasion assays performed immediately following tumour cell isolation. The relatively small biological sample sizes and limited availability of archived tumour sections represent important limitations of this study, particularly for the exploratory dissemination experiments and more detailed immune cell characterisation. The available tissue was insufficient for comprehensive analysis of additional immune cell populations and their activation states. Larger experimental cohorts, broader immune profiling and functional studies will be necessary to establish the mechanisms linking tumour induction protocols to subsequent disease outcomes, particularly tumour-derived signals, distant organ remodelling and disseminated tumour cell fate.

These limitations, nonetheless demonstrate that tumour induction protocols profoundly shape early tumour-host interactions, tumour microenvironmental evolution, which are known to impact dissemination behaviour.

Recognising and leveraging this variability offers a powerful opportunity to refine preclinical models to better reflect specific stages and contexts of human breast cancer progression, by integrating parameters such as tumour density, hypoxia signatures or immune infiltration patterns with human TNBC transcriptomic or spacial datasets.

## Conclusion

This study demonstrates that orthotopic implantation of breast cancer cells creates intrinsic experimental biases that profoundly influence tumour microenvironment and immune context. Tumour induction density governs HIF-1α signalling, vascular characteristics, necrosis, proliferation, and inflammatory infiltration, shaping tumour biology in ways that are not captured by tumour size or growth kinetics alone. These findings emphasise that orthotopic tumours generated using different protocols do not represent equivalent disease states, even within the same model system. Careful consideration of tumour induction parameters is therefore critical for accurately modelling human breast cancer progression, interpreting experimental outcomes, and designing preclinical studies that meaningfully inform translational and therapeutic strategies.

## Supporting information

Supplementary Materia

## Acknowledgements

The authors would like to thank Prof. Penelope Ottewell, University of Sheffield, for her generous gift of the 4T1^GFPLuc^ cells used to develop this study, and all past and present members of the Branco group who, while not directly contributing to the study, provided insights through actively engaging in discussions during the project that ultimately improved this manuscript. This work was partially funded by Breast Cancer Now (Scientific Fellowship ref.: 2014MaySF275).

## Author contributions

*Conceptualization:* C.B, G.A.; *Methodology:* S.McC, Y.L., G.A., N.L., A.E.; *Validation:* S. McC, Y.L., N.L; *Formal analyses*: S.McC., Y.L., C.M.B; *Investigation*: SMcL, Y.L, G.A.; *Resources*: C.B; *Writing – original draft preparation*: S.McC, C.M.B; *Writing – review & editing*: S. McC., Y.L., G.A., N.L., A.E., C.B; *Visualization*: S.McC., Y.L., C.M.B; *Supervision*: C.B; *Project administration*: C.M.B; *Funding Acquisition*: C.M.B

## Notes

### Competing Interest Statement

The authors have declared no competing interest.

