## Supplementary Materia for "ORTHOTOPIC TUMOUR INDUCTION PROTOCOL SHAPES TUMOUR MICROENVIRONMENT"

### *Supplementary material*

#### Figure S1

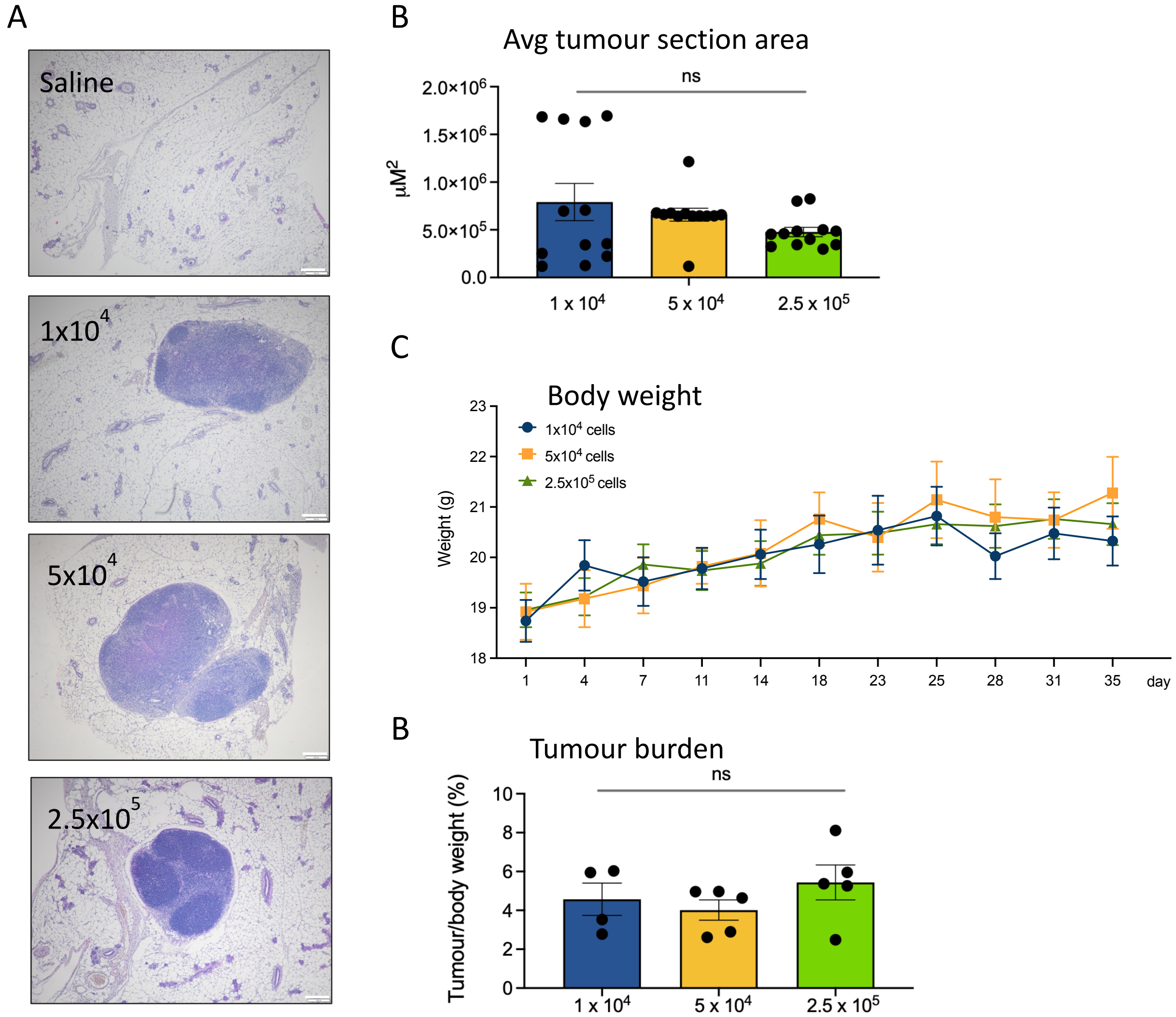


***Supplementary figure S1: Tumour induction protocol does not significantly affect tumour growth kinetics.*** *(A) Representative images at 4 x magnification of H&E stained mammary tumour sections from the different experimental groups, shown alongside a section from tumour-free mammary gland (saline, sham implantation); (B) Tumour section area for each experimental group was calculated for sections of 3 d tumours using FIJI; shown as average ± SEM. Significance was assessed by two-way ANOVA; n= 3 tumours per experimental group x 4 sections/tumour n=12 datapoints total per experimental group; (C) body weight records throughout experiment are shown as Avg ± SD; (D) tumour burden (tumour weight/body weight x 100) shown as Avg ± SEM*

### Figure S2

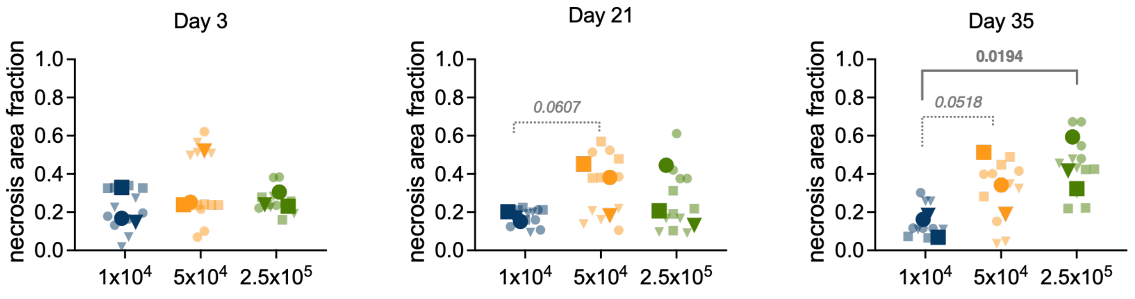


***Supplementary figure S2:*** *Average area of necrosis in primary tumours was calculated using QPath and plotted as a function of number of cells used to generate tumours (same data shown in Fig.2). Faded symbols represent one section of the tumour, solid symbols of same shape represent average for each tumour; Statistical significance was assessed by nested two-way ANOVA and p values showed when significant or approaching significance (italicised).*

**Figure S3**

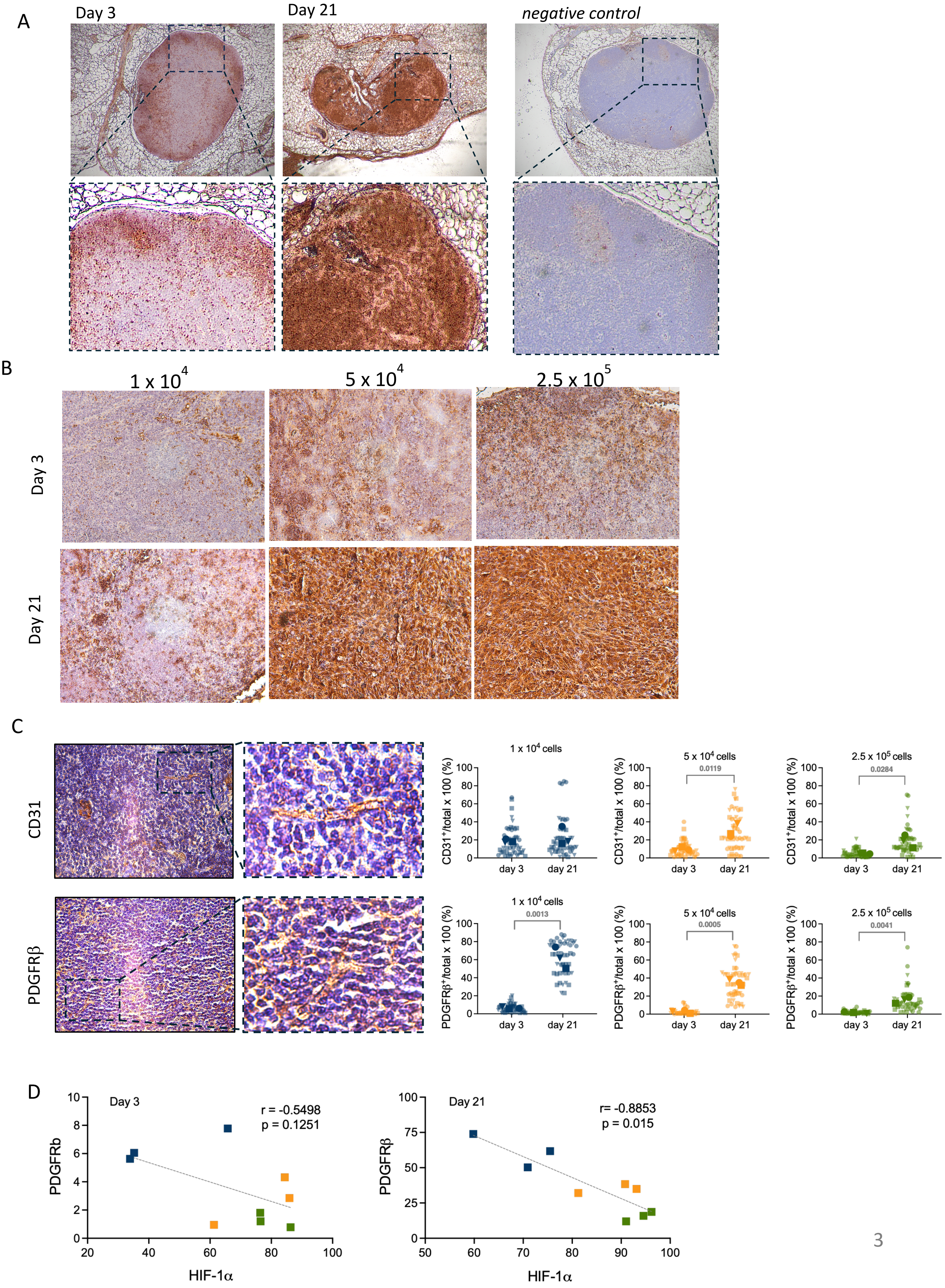
A
Day 3
Day 21
negative control
B
1 x 10^4
5 x 10^4
2.5 x 10^5
Day 3
Day 21
C
CD31
PDGFRβ
1 x 10^4 cells
5 x 10^4 cells
2.5 x 10^5 cells
0.0119
0.0284
0.0013
0.0005
0.0041
CD31+/total x 100 (%)
PDGFRβ+/total x 100 (%)
day 3
day 21
D
Day 3
r = -0.5498
p = 0.1251
PDGFRb
HIF-1α
Day 21
r= -0.8853
p = 0.015
PDGFRβ
HIF-1α

***Supplementary figure S3:*** *(A) Representative images of low magnification (4x) images of tumours at 3d and 21d post implantation stained for HIF-1$\alpha$ (5 x $10^5$ group shown, alongside negative control, digital amplification of selected areas shown under original image); (B) representative sections imaged at 10x of sections stained for HIF-1$\alpha$, and used for quantification for each time point and experimental group (data shown in shown in Fig. 3A); (C) Representative images of sections stained with endothelial cell marker CD31, and tumour pericyte marker PDGFR$\beta$ (digital amplification of identified areas are shown to the right of original image, obtained at 10x magnification); signal quantification (positively stained cells/total number of cells x 100) is plotted for each experimental group (blue, 1 x $10^4$; yellow 5 x $10^4$; green 2.5 x $10^5$) to show change over time; same data presented in Fig. 3B-C. Statistical significance was assessed by nested students t test and p values showed for significant or near significant (italicised) differences. D. Relationship between HIF-1α and PDGFRβ staining in tumours collected 3 and 21 days post implantation. Each point represents the average staining measurement from one biological replicate (mouse), with colours indicating the implantation groups (blue, 1 × $10^4$; yellow, 5 × $10^4$; green, 2.5 × $10^5$ cells). Pearson correlation was calculated using paired tumour-level measurements (n = 9 tumours per time point). Regression lines are shown for visualisation.*

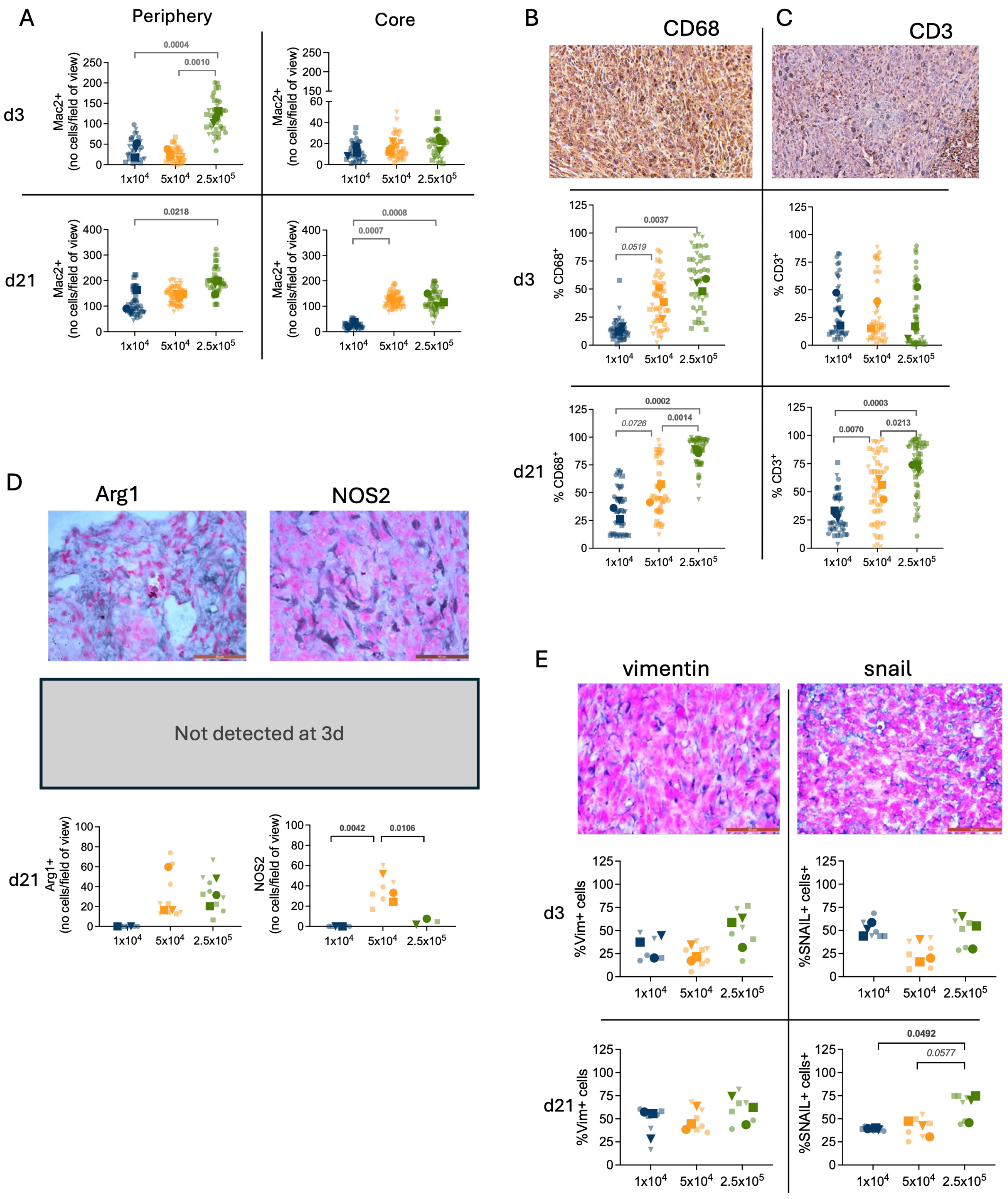


***Supplementary figure S4.*** *(A) The number of MAC2+ cells per field of view (fov) of peripheral and core macrophages shown in Fig. 4 are presented for each time point to compare fraction between experimental groups (instead of within each group over time); (B) Representative images of CD68 stained sections and quantification in each experimental cohort and time-point (no. CD68+/total no. cells x 100); (C) representative images and quantification of CD3 cells (no.CD3+/total no. cells x 100), as in (B); (D) Representative images of macrophage polarization markers Arg 1 (M2, left) and NOS2 (M1, right) and quantification of signal from sections of 21d tumours (no signal detected in sections of 3d tumours); (E) Representative markers of mesenchymal status Vimentin (left) and EMT-associated transcription factor SNAIL (right) are shown, with quantification for each*

*time point and experimental group. Matching faded symbols represent readouts of sections from the same biological replicate (tumour); average for each tumour is shown with identical symbols in solid colour; n=3 tumours, ≥ 4 sections stained per tumour, ≥ 4 random images obtained for quantification (minimum 48 datapoints per tumour); Statistical significance was determined using nested two-way ANOVA, and p values shown for significant or near significant (italicised) differences.*

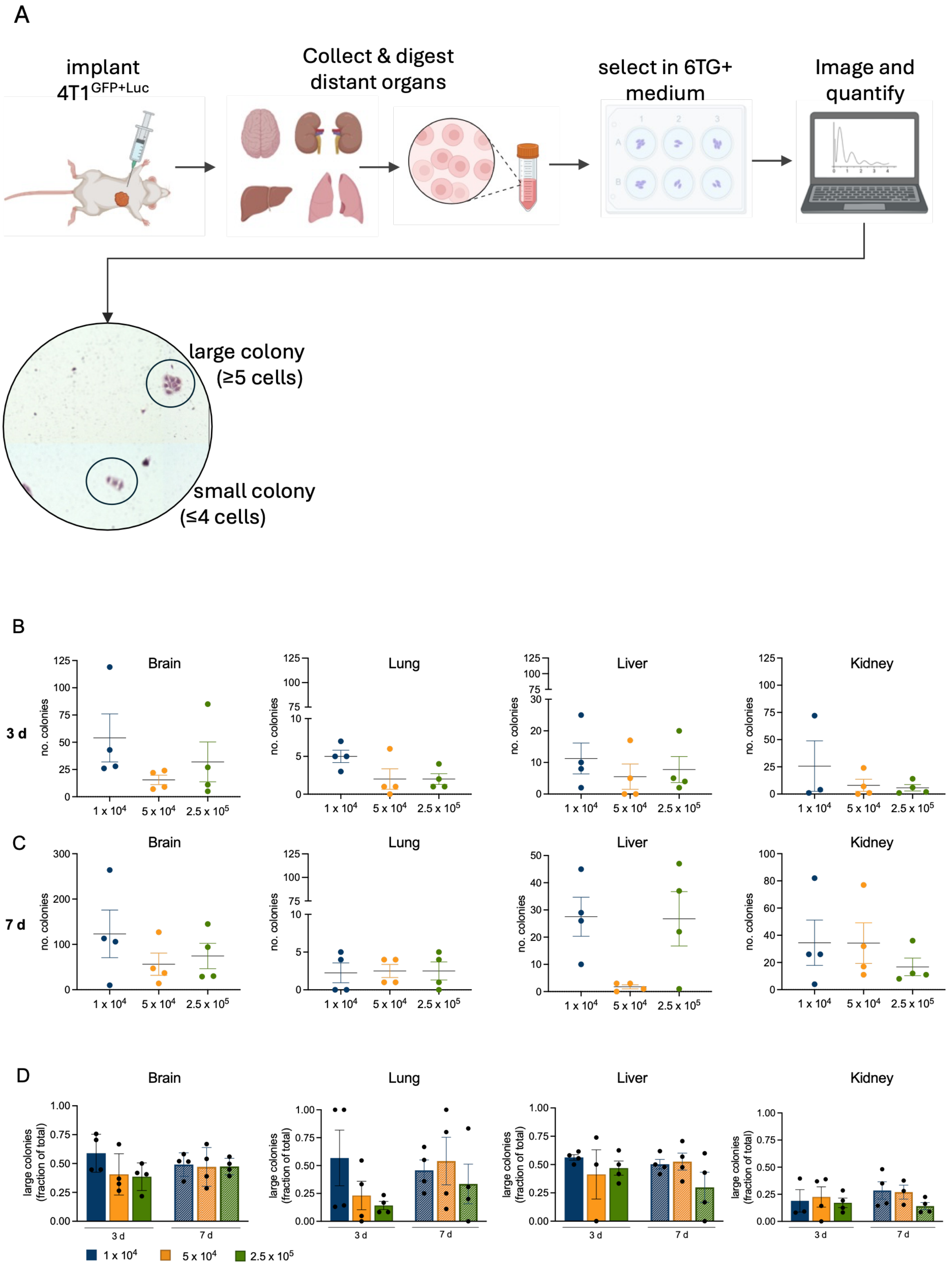


***Supplementary figure S5.*** **Tumour cell dissemination is a function of primary tumour density.** (A) Visceral organs to represent metastasis prone (lung, brain and liver) and metastasis resilient (kidney) organs were excised 3 and 7 d post implantation of mammary tumours (generated with 1 x $10^4$, 5 x $10^4$, or 2.5 x $10^5$), each digested into single cell suspensions, then plated 6-thioguanine containing medium to select 4T1 cells. Stained colonies were imaged 4 d later for quantification; representative image of 4T1 colonies are shown; (B) Number of total colonies obtained per organ when extracted 3d (B) or 7 d (C) post implantation of primary tumour. (D) Fraction of large colonies (≥ 5 cells) from total number of colonies found in each organ after 3 d and 7 d post implantation. N=4 mice per cohort; No statistically significant differences were seen between experimental groups (assessed by 2-way ANOVA).

Supplementary Table 1. **List of Reagents**

| Deparaffinisation and staining | | | | |
|---|---|---|---|---|
| Reagent | Supplier | Catalogue Number | Working solution | Application |
| Histoclear | Sigma | H2779-1L | As supplied | Deparaffinisation |
| Harris Haematoxylin | Leica | 3801560 | As supplied | Nuclear stain |
| Haematoxylin (alternative) | Sigma | GHS232 / 1051750500 | As supplied | Nuclear stain |
| $NaHCO_3$ | VWR | 27775.293 | 0.1%, pH 8 | Bluing Agent |
| Eosin | Sigma | 1.09844.1000 / 1098441000 | With acetic acid added at 1:500 | Cytoplasm stain |
| Ethanol, pure | Sigma-Aldrich | E7023 | 50%, 70%, 95%, 100% in di$H_2O$ | Dehydration |
| Quick Hardening Medium (Eukitt) | Sigma / Fluka | 03989 / BCBT0137 | As supplied | Mounting |
| Bovine Gelatin | Sigma | G9391 | Glycerol jelly mounting medium: 5g Gelatin, 50 mL $H_2O$, 50 mL Glycerol | |
| | Sigma | G5516 | | |
| **Antigen retrieval and blocking** | | | | |
| Reagent | Supplier | Cat. No. | Working solution | Application |
| Antigen Retrieval Buffer (Citrate) | Vector | H-3300 | 1:100 in dd$H_2O$ | Retrieval |
| $H_2O_2$ | Aldrich / Sigma | 216763 | 3% in Methanol | Peroxidase quenching |
| Universal Protein Blocking Reagent | GeneTex | GTX30693 | – | Protein block |
| Goat Serum | Sigma | G9023 | 5% in PBS | Blocking |
| Donkey Serum | Sigma | D9663 | 5% in PBS | Blocking |
| Biotin/Streptavidin Block | Vector Labs | SP2002 | 1 drop each in 5mL PBS | Blocking |
| MOM Blocking Solution | Vector | MKB-2213 | – | Blocking |
| Triton X-100 | | | | |
| **Primary antibodies** | | | | |
| Target | supplier | Cat. number | Dilution used | Host |
| SNAIL | Novus Biologicals | NBP2-27293 | 1:200 | Rabbit |
| Vimentin | Novus Biologicals | NB300-223 | 1:500 | Rabbit |
| Arg1 | Santa Cruz Biotech | sc-20150 | 1:200 | Rabbit |
| NOS2 | Santa Cruz Biotech | sc-651 | 1:200 | Rabbit |
| CD3 | Novus Biologicals | ab5690 | 1:75 | Rabbit |

| | | | | |
|---|---|---|---|---|
| PCNA | Novus Biologicals | NB500-106 | 1:100 | Mouse |
| CD68 | Abcam | ab125212 | 1:100 | Rabbit |
| CD31 | Abcam | ab28364 | 1:100 | Rabbit |
| PDGFRβ | R&D Systems | AF1042 | 1:100 | Goat |
| HIF1α | Novus Biologicals | NB100-105 | 1:100 | Mouse |
| Mac2 | Biolegend | BIOLOGEND 125403 | 1:400 | Rat |
| Ki67 | R&D systems | AF7649 | 1:150 | Sheep |
| **Secondary Antibodies** | | | | |
| Host and Target | Supplier | Cat. No. | Dilution used | Conjugate |
| Goat Anti-Chicken | Santa Cruz Biotech | sc-2428 | 1:200 | HRP |
| Goat Anti-Rabbit | Vector | BA-1000 | 1:200 | Biotin |
| Goat Anti-Mouse | Invitrogen | D20691 | 1:200 | Biotin |
| Donkey Anti-Goat | Santa Cruz Biotech | sc-2042 | 1:200 | Biotin |
| Goat anti-rat | Santa Cruz Biotech | Sc-2041 | 1:200 | HRP |
| Donkey anti-sheep | Abcam | Ab6900 | 1:200 | HRP |
| **Detection Reagents** | | | | |
| Reagent | Supplier | Cat. No. | Working solution | Application |
| Streptavidin-HRP | Abcam | ab64269 | Per manufacturers instructions | Signal amplification |
| ExtrAvidin Peroxidase | Sigma | E2886 | 1:100 in PBS | Signal amplification |
| HRP Substrate SG | Vector SG | SK-4700 | 3 drops each in 5 mL PBS | Chromogen |
| DAB Substrate | Abcam | ab64238 | Per manufacturers instructions | Chromogen |
| Nuclear Fast Red Powder | Sigma-Aldrich | 229113 | 0.1 g in 250mL $H_2O$ + 5 g Alum Sulphate for nuclear staining | |
| Aluminium sulphate | Sigma-Aldrich | 368458 | | |
| Crystal Violet | Alfa Easer | 022866-22 | in 70% ethanol | Cell staining |
| **Cell Culture** | | | | |
| **Reagent** | **Supplier** | **Cat. No.** | **Dilution** | **Application** |
| DMEM | Gibco | 41965-039 | (n/a) | Growth/Digestion Medium |
| Collagenase A | Roche | 11088793001 | 0.2% in PBS | Tissue digestion |
| ACK Lysis Buffer | Gibco | A1049201 | (n/a) | RBC lysis |
| DNase I | Roche | 10104159001 | 10 μg/mL (Brain) | Tissue digestion |

| | | | | |
|---|---|---|---|---|
| BSA | Sigma | A7906 | 20% in DMEM | Myelin removal |
| RPMI | Gibco | 21875-034 | (n/a) | Media |
| FBS | Gibco | 10270-106 | 10% in RPMI | Supplement |
| 6-Thioguanine (6-TG) | ThermoFisher Scientific | B21280.06 | 60 mM | Selection agent |
